# Functional Assays Reclassify Suspected Splice-Altering Variants of Uncertain Significance in Mendelian Channelopathies

**DOI:** 10.1101/2022.03.14.484344

**Authors:** Matthew J. O’Neill, Yuko Wada, Lynn D. Hall, Devyn W. Mitchell, Andrew M. Glazer, Dan M. Roden

## Abstract

**Background:** Rare protein-altering variants in *SCN5A, KCNQ1*, and *KCNH2* are major causes of Brugada Syndrome (BrS) and the congenital Long QT Syndrome (LQTS). While splice-altering variants lying outside 2-bp canonical splice sites can cause these diseases, their role remains poorly described.

**Objective:** We implemented two functional assays to assess 12 recently reported putative splice-altering variants of uncertain significance (VUS) and 1 likely pathogenic (LP) variant without functional data observed in BrS and LQTS probands.

**Methods:** We deployed minigene assays to assess the splicing consequences of 10 variants. Three variants incompatible with the minigene approach were introduced into control induced pluripotent stem cells (iPSCs) by CRISPR genome editing. We differentiated cells into iPSC-derived cardiomyocytes (iPSC-CMs) and studied splicing outcomes by reverse transcription-polymerase chain reaction (RT-PCR). We used the American College of Medical Genetics and Genomics functional assay criteria (PS3/BS3) to reclassify variants.

**Results:** We identified aberrant splicing, with presumed disruption of protein sequence, in 8/10 variants studied using the minigene assay and 1/3 studied in iPSC-CMs. We reclassified 9 VUS to LP, 1 VUS to Likely Benign, and 1 LP variant to pathogenic.

**Conclusions:** Functional assays reclassified splice-altering variants outside canonical splice sites in BrS- and LQTS-associated genes.

## Introduction

The arrhythmia syndromes Brugada Syndrome (BrS) and congenital Long QT Syndrome (LQTS) are rare autosomal dominant Mendelian diseases, mainly involving variants in cardiac ion channels. BrS is associated with loss-of-function (LoF) variants in the *SCN5A* sodium channel gene in 20% of patients, while LoF variants in the potassium channel genes *KCNQ1* and *KCNH2* and gain-of-function (GoF) variants in *SCN5A* are found in 80% of patients with congenital LQTS ^1,2^. These diseases contribute to the >250,000 cases of sudden cardiac death (SCD) each year in the US through fatal ventricular arrhythmias. When affected heterozygotes are recognized early, BrS and LQTS can be clinically managed through medications and/or interventional therapies. However, when not recognized through medical or genetic screening, SCD may present as the sentinel disease manifestation^3^. Genetic sequencing therefore offers the opportunity to uncover risk in 1) clinically unrecognized heterozygotes identified in unascertained population sequencing studies, or 2) family members of a recognized proband. Multiple approaches are in development to support genome-first approaches and cascade screening, but these efforts are still hampered by issues of clinical interpretability^4,5^. The American College of Medical Genetics and Genomics (ACMG) has provided a framework for variant interpretation, spanning benign (B) to pathogenic (P) based on criteria including variant functional evidence, population minor allele frequencies, segregation within families, and computational predictions, among others^6^. While panel sequencing and whole exome/genome sequencing are improving our broader understanding of the genetic basis of disease, a large proportion of detected variants are classified as Variants of Uncertain Significance (VUS) and are therefore not clinically actionable^7^.

Protein-altering variants (including nonsense, frameshift, and missense variants) are a major focus of clinical attention. Variants affecting RNA splicing occupy a comparatively less explored fraction of the genome, despite estimates that they contribute to up to 10% of pathogenic variants in Mendelian diseases^8^. Aberrant splicing from rare variants that alter the 2-bp canonical splice sites flanking each exon are typically characterized as meeting the PVS1 criterion, and therefore often are classified as P/LP. However, rare splice-altering variants falling outside of these 2-bp sites are more difficult to interpret. Aberrant splicing may arise from more distant intronic or exonic variation by introducing or ablating splice acceptors or donor sequences or by affecting regulatory splicing enhancers or silencers (Figures 1A and 1B)^9^. *In silico* predictors of splicing consequences have historically had only modest predictive ability.^10,11^ However, recent tools leveraging advances in machine learning and large RNA-seq datasets raise the possibility that *in silico* splicing predictors may be increasingly used to facilitate interpretation of suspected splice-altering variation^8^.

**Figure 1.**
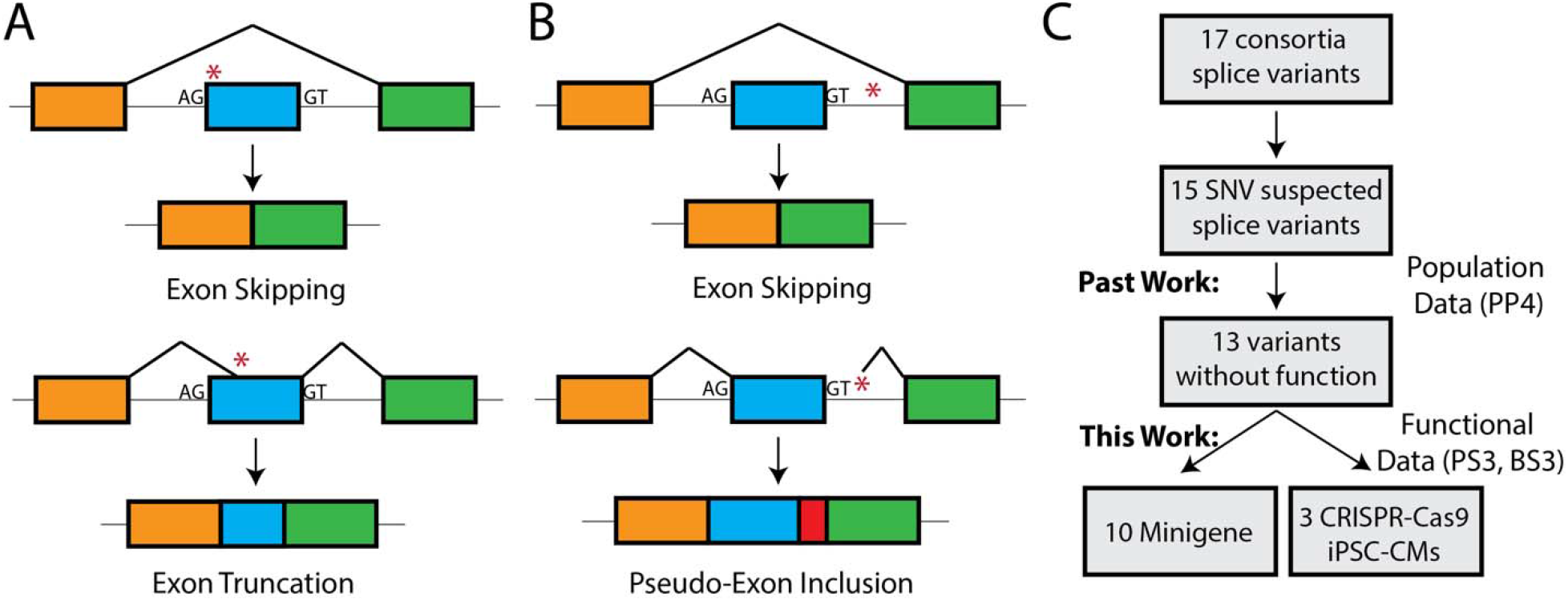
Aberrant splicing from *cis-*genetic variation and previous variant prioritization. A) Variants in an exon near the 2-bp splice acceptor (AG) or donor (GT) can disrupt the recognition of the canonical site and lead to exon skipping. Alternatively, variants within the exon may introduce a new splice acceptor or donor site and lead to truncation of the exon, which may introduce frameshifts or damage protein function. This schematic features canonical AG-GT splice sites; rare alternate splice sites are possible. B) Intronic variants adjacent to the 2-bp splice acceptor or donor may disrupt spliceosome recognition of the canonical splice site and lead to exon skipping. Deeper intronic variants may introduce cryptic splice acceptor or donor sites that may lead to transcription of intronic regions (pseudo-exons) that may disrupt the reading frame or compromise protein function. This schematic features canonical AG-GT splice sites; rare alternate splice sites are possible. C) A recent variant interpretation effort curated LQTS and BrS heterozygotes and reported variants. 17 putative splice-altering variants were reported, of which 15 were SNV. Of these, 13 were previously not functionally characterized. We analyzed 10 of 13 using a standard minigene approach and 3 using RT-PCR assays from iPSC-CMs.

In this study, we used minigene assays in Human Embryonic Kidney 293 (HEK293) cells to study variant consequences on splicing for 10 putative splice-altering variants in arrhythmia genes identified in a recent cohort of BrS and LQTS patients (Figure 1C). For three variants incompatible with minigene assays, we examined the impact on splicing in induced pluripotent stem cell-derived cardiomyocytes (iPSC-CMs) that were edited with CRISPR-Cas9^12^. We applied these functional splicing assays to reclassify a total of 11 putative splice-altering variants within the ACMG framework, including 9 VUS to Likely Pathogenic.

## Methods

### 1. General Methods

#### 1.1 Selection of Variants

Putative splice-altering variants were identified from a recent curation of variants from Walsh et al.^13^ Variants were considered putative splice-altering variants if they were identified in at least 1 case of BrS (*SCN5A*) or LQTS (*KCNQ1* and *KCNH2*), and were not predicted to affect the coding sequence of the protein. Additionally, all candidate variants had an allele frequency lower than 2.5e-5 in the Genome Aggregation Database (gnomAD)^7^, a cutoff derived from theoretical predictions and empirical measurements of maximum allele frequency for Mendelian arrhythmia variants^13,14^. Variant case^13^ and gnomAD^7^ counts are shown in Table 1. We used the following transcripts throughout our study: ENST00000155840 (*KCNQ1*), ENST00000262186 (*KCNH2*), and ENST00000333535 (*SCN5A*).

**Table 1.** Summary of variants with aggregate SpliceAI scores, assay results, case and control frequencies, and ACMG reclassifications.

| Gene | Variant | Aggregate SpliceAI | Assay Type | Assay Result | Case / gnomAD | ACMG Post-Assay | Reclassification |
| --- | --- | --- | --- | --- | --- | --- | --- |
| *SCN5A* | c.393-5C>T | 0.21 | iPSC-CM | Normal | 1 / 0 | PM2, BS3, BP4 | VUS → LB |
| *SCN5A* | c.1890G>A | 0.72 | Minigene | Aberrant | 1 / 0 | PM2, PS3, | VUS → LP |
| *SCN5A* | c.4299+6T>C | 0.75 | Minigene | N/A | 1 / 0 | PM2 | VUS → VUS |
| *SCN5A* | c.4299G>C | 0.80 | Minigene | N/A | 1 / 0 | PM2 | VUS → VUS |
| *SCN5A* | c.4437+5G>A | 0.69 | iPSC-CM | Aberrant | 2 / 0 | PM2, PS3 | VUS → LP |
| *SCN5A* | c.4719C>T | 0.96 | Minigene | Aberrant | 2 / 2 | PM2, PS3,<br>PS4_strong, PP3 | LP → P |
| *KCNH2* | c.1128G>A | 0.34 | Minigene | Aberrant | 2 / 0 | PM2, PS3 | VUS → LP |
| *KCNH2* | c.2145G>A | 0.25 | Minigene | Aberrant | 2 / 0 | PM2, PS3, BP4 | VUS → LP |
| *KCNH2* | c.2398+5G>T | 0.45 | Minigene | Aberrant | 2 / 0 | PM2, PS3 | VUS → LP |
| *KCNQ1* | c.386+6T>G | 0.96 | iPSC-CM | Benign | 1 / 0 | PM2, PS3, PP3 | VUS → LP |
| *KCNQ1* | c.477+5G>A | 0.75 | Minigene | Aberrant | 7 / 3 | PM2, PS3,<br>PS4_strong | LP → P |
| *KCNQ1* | c.683+5G>A | 0.29 | Minigene | Aberrant | 3 / 4 | PM2, PS3,<br>PS4_moderate | VUS → LP |
| *KCNQ1* | c.1032+5G>A | 0.95 | Minigene | Aberrant | 2 / 0 | PM2, PS3, PP3 | VUS → LP |
| *KCNQ1* | c.1032G>T | 0.74 | Minigene | Aberrant | 1 / 0 | PM2, PS3 | VUS → LP |
| *KCNQ1* | c.1032G>A | 0.68 | Minigene | Aberrant | 9 / 0 | PM2, PS3,<br>PS4_strong | LP → P |

#### 1.2 Interpretation of Functional Assays

For minigene assays, we performed triplicate amplification of RT-PCR products and quantified the fraction of Percent Spliced In (PSI) of the wild-type (WT) exon cassette from gel intensities using ImageJ^15^. All bands were confirmed by Sanger sequencing. We deemed minigene results inconclusive if the PSI of WT plasmid was less than 30%. We considered a >50% disruption of WT splicing to fulfill the PS3 criteria, while the absence of splice perturbation (<10% change in PSI compared to WT) fulfilled the BS3 criteria. Statistical analyses were performed with a two-tailed unpaired t-test implemented in R. Error bars correspond to the standard error of the mean.

### 2. Minigene Assay

#### 2.1 Minigene constructs

A schematic of the minigene construct is shown in Figure 2A. PCR products bearing the exon of interest surrounded by 100 nucleotides of intronic DNA were amplified from healthy control gDNA (primers in Table S1). PCR products were gel extracted (Qiagen) then restriction digested with SalI and NotI (New England Biolabs). The plasmid pET01 (MoBiTec GmbH, Göttingen, Germany) similarly underwent double digestion with SalI and NotI, followed by treatment with Calf Intestinal Phosphatase (Promega). The digested products were ligated together with T4 ligase (New England Biolabs). An aliquot of the ligation mix was transformed into DH5α competent cells by heat shock (Thermo Scientific) and cells were grown overnight at 37°C incubated with LB broth and 1 mg/mL ampicillin. Plasmid DNA was extracted using the Qiagen Miniprep kit. Each variant of interest was introduced using the QuikChange Lightning Multi Kit (Agilent) with 1 primer per variant using primers designed using the online QuikChange Primer Design tool. All primers used in this study are presented in Tables S1. The sequences of wild-type and variant plasmids were confirmed using Sanger sequencing and the entire insert sequences were compared to the GrCh38 reference genome sequence to ensure there were no unanticipated variants present.

**Figure 2.**
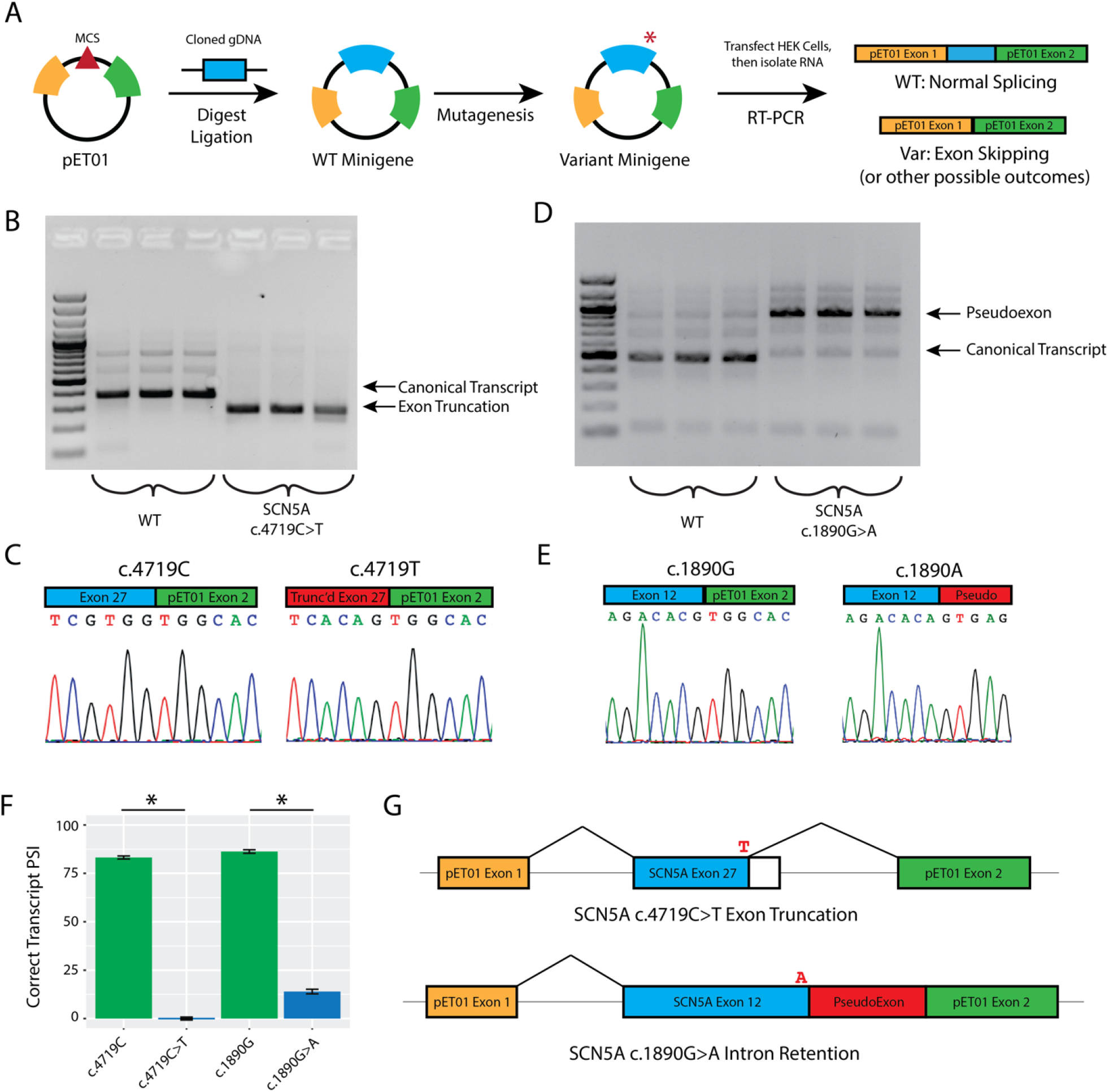
Minigene assay and studies on *SCN5A*. A) Schematic of minigene assay. The PCR product of an exon and segments of flanking introns of interest is cloned into a multiple cloning site (MCS) between two known exons. Mutagenesis inserts the variant. After transfection of the WT and variant plasmids into HEK cells, RNA is isolated, and RT-PCR provides cDNA for analysis of transcript composition. B) Gel of PCR products for the known exon truncating variant c.4719C>T and WT plasmid. The WT band is consistent with higher molecular weight compared to the truncated form induced by the aberrant splice donor gain. All minigene assays are presented in triplicate. C) Sanger sequencing of the exon-exon junctions of WT and truncated RT-PCR products. D) Gel of the WT exon 12 and c.1890G>A RT-PCR products implicating a pseudo-exon gain induced by the variant. E) Confirmatory Sanger sequencing of RT-PCR products for the WT and c.1890G>A VUS. F) Quantification of WT and variant Percent Spliced In (PSI). * indicates p < 0.01, two-tailed t-test. G) Schematic of assay results. *SCN5A* c.4719C>T truncates exon in our assay, whereas the *SCN5A* variant c.1890G>A induces intron retention.

#### 2.2 Cell Culture and Transfection

HEK cells were cultured at 37°C in humidified 95% air/5% CO_2_ incubator in “HEK media”: Dulbecco’s Eagle’s medium supplemented with 10% fetal bovine serum, 1% non-essential amino acids, and 1% penicillin/streptomycin. At 30% confluency, HEK cells were transfected with 500 ng of plasmid using FuGENE 6 (Promega) following manufacturer’s instructions.

#### 2.3 RNA Isolation and Reverse Transcription-Polymerase Chain Reaction (RT-PCR)

48 hours after transfection, HEK cells were washed with PBS, treated with Trypsin for 2 minutes, and harvested with HEK media. Following centrifugation at 300g for 5 minutes, the supernatant was aspirated, and the pellet was immediately placed on ice. RNA was extracted from the pellet using the Qiagen RNeasy Minikit. Reverse transcription was performed using SuperScript® III System (Invitrogen) using gene specific primers (Table S1). PCR was performed using GoTaq PCR Master Mix (Promega) with primers mo37 and ag489. For exon skipping events, the primer mo102 was used to confirm the splice junction by Sanger sequencing. DNA was amplified using a touchdown protocol: 98°C for 30”, (98°C for 10s, 65°C for 30s [decreasing 1°C/cycle], 72°C for 60s) x 10 cycles, (98°C for 10s, 55°C for 15s, 72°C for 60s) x 30 cycles.

#### 2.4 Gel Electrophoresis and Extraction of RT-PCR Products

RT-PCR products were separated by gel electrophoresis using 2% agarose gels in TAE buffer. Bands were visualized with UV light and excised followed by DNA isolation (Qiagen Gel Extraction kit). Extracted DNA was analyzed by Sanger sequencing.

### 3. Analysis of genome-edited iPSC-CMs

#### 3.1 CRISPR-Cas9

Primers for CRISPR gRNAs were designed using the online CRISPOR tool^16^. Guides were cloned into SpCas9-2A-GFP (pX458)^17^ plasmid (Addgene #48138, a gift of Feng Zhang). CRISPR guides used in this study are presented in Table S1. The guide plasmid and a 150 bp single-stranded homology directed repair oligonucleotide template with the variant of interest and a PAM site variant to disrupt re-cutting were co-electroporated into control iPSCs from a healthy donor using a Neon Transfection System (ThermoFisher MPK5000).^18^ After 48 h, GFP+ single cells were sorted on a BD Fortessa 5-laser instrument and sub-cloned. Each sub-clone was genotyped by variant-specific primers. Sanger sequencing of amplified DNA identified clones with the correct heterozygous variant (Table S1).

#### 3.2 Cardiac Differentiation of iPSC

iPSCs were maintained on plates coated with Matrigel (BD Biosciences) in mTeSR plus media (STEMCELL) at 37°C in a humidified 95% air/5% CO_2_ incubator. At 60-80% confluency, iPSCs were differentiated into iPSC-CMs using a monolayer chemical method as previously described^19^.

#### 3.3 RNA isolation and RT-PCR

iPSC-CMs were harvested at day 30 of differentiation following the RNeasy Minikit protocol (Qiagen). RT-PCR was performed as above with variant-specific primers for each step (Table S1). The amplified cDNA was separated using 2% agarose gel electrophoresis, extracted as above, and analyzed with Sanger sequencing. For TA cloning, the gel extracted product was ligated into the pGEM®-T Vector (Promega) using the supplied kit per manufacturer’s instructions. After ligation and transformation, single colonies were grown up, DNA was extracted with the Qiagen Miniprep kit as above, and transcript composition determined by Sanger sequencing.

### 4. SpliceAI

*SpliceAI*. The *in silico* predictor SpliceAI was used to predict the splicing consequences of each assayed variant. Variants were manually interrogated using the web-based interface hosted at https://spliceailookup.broadinstitute.org/ using GRCh38 coordinates and a maximum distance of 1000 bp. For each variant, four predicted probabilities were obtained: probabilities of introducing a splice Acceptor Gain (AG), Acceptor Loss (AL), Donor Gain (DG), or Donor Loss (DL) accompanied by a predicted position of such change relative to each variant. We computed an aggregate SpliceAI score to incorporate contributions from each of the 4 predicted categories using a previously described formula^20^:

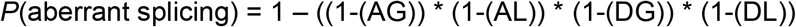

We considered a high likelihood of disrupting splicing to be greater than 0.80, and a low likelihood less than 0.25.

### 5. ACMG Reclassification

There has been extensive use and validation of this minigene assay in our study and previous studies^21-23^ and there is a clear link between heterozygous LoF variants of the 3 studied genes with Long QT Syndrome (*KCNQ1* and *KCNH2*) or Brugada Syndrome (*SCN5A*)^1,2^. Therefore, we considered a conclusive assay result (as defined above) that resulted in >50% disruption of splicing to fulfill the PS3 criterion, while the absence of splice perturbation fulfilled the BS3 criterion. We applied the PP3 criterion (computational prediction shows deleterious effect) to variants with an aggregate SpliceAI score >0.8 and the BP4 criterion (computational prediction shows no deleterious effect) to variants with an aggregate SpliceAI score <0.25. We supplemented the previously published ACMG criteria from Walsh et al. (Supplemental Table 3 and Table 1) with these functional assays and computational findings for reclassification, using the University of Maryland ACMG online tool to implement the criteria^13,24^.

## Results

### Minigene Assay Reveals Aberrant Splicing

We used a minigene construct previously deployed by our group^21^ and others^25,26^ to study the effects of *SCN5A* potential splice-altering variants that have been observed in one or more patients with BrS (Figure 2A). This assay allows the direct comparison of WT versus variant splicing outcomes by studying a specific exon and flanking intronic regions of interest. We first studied the known likely pathogenic (LP) exon-truncating *SCN5A* synonymous variant c.4719C>T in our system, which had previously been determined to disrupt splicing in a minigene assay^27^. Compared to the respective3 WT minigene construct, the size of the RT-PCR product from the c.4719C>T minigene was consistent with cDNA truncation (Figure 2B). This was confirmed by Sanger sequencing (Figure 2C). We next studied the *SCN5A* VUS c.1890G>A and its WT counterpart. In the minigene assay, c.1890G>A resulted in retention of intron 10 by a splice donor loss as evidenced by RT-PCR product gel band size and confirmatory Sanger sequencing (Figure 2D and 2E). Quantification of the gel band intensity showed significant changes in the WT exon cassette percent spliced in (PSI) in both cases (Figure 2F). The effects on transcript composition of these two variants are shown schematically in Figure 2G. This approach was attempted for two additional *SCN5A* VUS, c.4299G>C and c.4299+6T>C, but the results of the minigene assay were inconclusive due to poor splicing in of the WT exon (Supplementary Figure 1).

We studied three *KCNH2* VUS c.1128G>A, c.2145G>A, and c.2398+5G>T discovered in the LQTS cohort^13^. The variant c.1128G>A induced exon skipping, which is predicted to disrupt the downstream reading frame based on exon size (Figure S2). The variant c.2145G>A led to an exon skipping event at exon 8 (Figure S2), similarly resulting in a predicted frameshift. The intronic variant c.2398+5G>T behaved similarly, leading to exon skipping (Figure S2). This exon 9 skipping event is also predicted to lead to a frameshift. Quantification of the respective WT exon cassette PSI showed significant changes in all cases (p < 0.01, two-tailed t-test) (Figure 3A). Schematic depictions of the major splicing outcomes on the transcript are shown in Figure 3B. Raw gels and Sanger traces for all major splicing outcomes are shown in Supplementary Figure 2.

**Figure 3.**
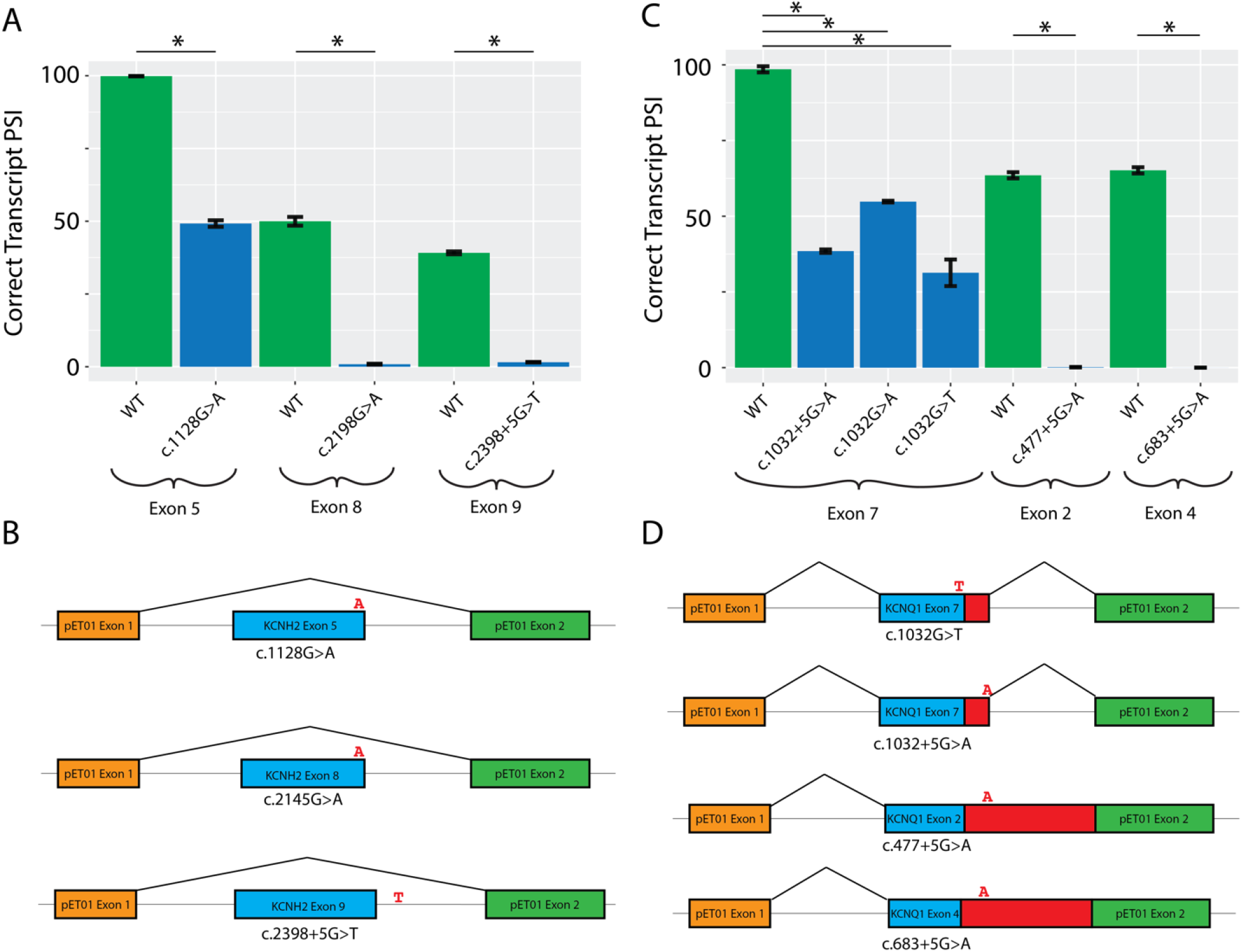
Minigene analysis of *KCNH2* and *KCNQ1* splice variants. A) Quantification of *KCNH2* band intensity corresponding to the PSI of the WT and variant bands (* indicates p < 0.01, two-tailed t-test). B) Schematic depiction of the splicing outcome associated with the highest intensity band for each *KCNH2* variant. C) Quantification of normal exon PSI across all samples (* indicates p < 0.01, two-tailed t-test). D) Schematic depictions of the major splicing outcome for each variant.

We studied 5 candidate LQTS splicing variants in *KCNQ1*. 3 *KCNQ1* VUS (c.683+5G>A, c.1032+5G>A, c.1032G>T) and 2 *KCNQ1* LP variants (c.477+5G>A and c.1032G>A) were studied. As a control, we first studied c.1032G>A, one of the most common causes of LQTS among Japanese probands (29 cases, 0 gnomAD controls).^28^ This variant has been previously shown to affect splicing *in vitro* and did so in our assay as well, leading to 2 alternative pseudoexon inclusions (Figure S3). The adjacent intronic VUS, c.1032+5G>A, had a similar gel electrophoresis profile and led to multiple pseudoexon inclusions with variable respective cryptic donor sites. A second VUS near this splice junction, c.1032G>T, was also functionally assayed. In this case, 3 pseudoexons and an exon skipping event were observed, with a small amount of WT transcript overlapping with a pseudoexon (Figure S3). These consequences are predicted to have similarly deleterious effects on the as those previously studied for c.1032G>A^28^. These results show that the *KCNQ1* exon 9 splice donor site is highly intolerant to variation. The previously unstudied LP variant, c.477+5G>A, was assayed and showed nearly complete loss of WT splicing, with inclusion of 2 unambiguously defined pseudoexons (Figure S3). The retained intron sequence contains a stop codon that would disrupt protein function. In our functional studies, introduction of the VUS c.683+5G>A variant completely abrogated WT splicing, while also introducing a pseudoexon and retaining an exon skipping event moderately observed in the WT (Figure S3). The retained intronic sequence would also lead to a stop codon inclusion, which is predicted to disrupt protein function. Quantifications of splicing outcomes for the WT PSI are shown in Figure 3C (p < 0.01 in all cases, 2-tailed t-test). Major splicing outcomes for each variant are depicted schematically in Figure 3D. Raw gels and Sanger traces for all major splicing outcomes are shown in Supplementary Figure 3.

### CRISPR/Cas9 and iPSC-CMs Reveal Aberrant Splicing

Although the minigene assay can reliably assess many variant effects on splicing, it is incompatible with assaying the first or last exon of a gene, or assaying rare splice sites that do not use the standard AG-GT sites^29^. Three candidate splice-altering VUSs were incompatible with minigene assays: *KCNQ1* c.386+6T>G (first exon), *SCN5A* c.393-5C>T (AC-GT splice site), and *SCN5A* c.4437+5G>A (AG-AT splice site). These three variants were therefore studied by introducing each variant with CRISPR-Cas9 into healthy control iPSCs, differentiating into iPSC-CMs, and then assessing splicing consequences by RT-PCR of isolated RNA^17^.

The RT-PCR products of the *SCN5A* VUS c.4437+5G>A and WT RNA were approximately the same size by gel electrophoresis (Figure 4A); however, Sanger sequencing of relevant exon junctions showed aberrant splicing of the edited allele (Figure 4B). We used negative and positive controls in all RT-PCR experiments corresponding to no PCR input and a WT *SCN5A* or *KCNQ1* cDNA construct. TA cloning and sequencing revealed 2 major transcripts corresponding to the WT and predominant edited allele product, inducing a frameshifting variant (Figure 4B). Splicing predictions suggested a cryptic donor site that would lead to a frameshift of the transcript, matching the experimentally observed transcript (Figure 4C). The variant *SCN5A* c.393-5C>T was previously studied by minigene assays and shown to induce exon skipping^30^; however, due to the non-standard splice site usage, this specific implementation of the assay may lead to inappropriate conclusions. In the CRISPR-Cas9 iPSC-CM model, we observed no change in splicing by RT-PCR (Figures 4D and 4E). RT-PCR of the *KCNQ1* VUS c.386+6T>G ALSO showed no disruption of splicing by gel electrophoresis or by Sanger sequencing (Figures 4F and 4G). Altogether, these studies implicated *SCN5A* c.4437+5G>A as splice-altering, and *SCN5A* c.393-5C>T and *KCNQ1* C.386+6T>G as not splice-altering.

**Figure 4.**
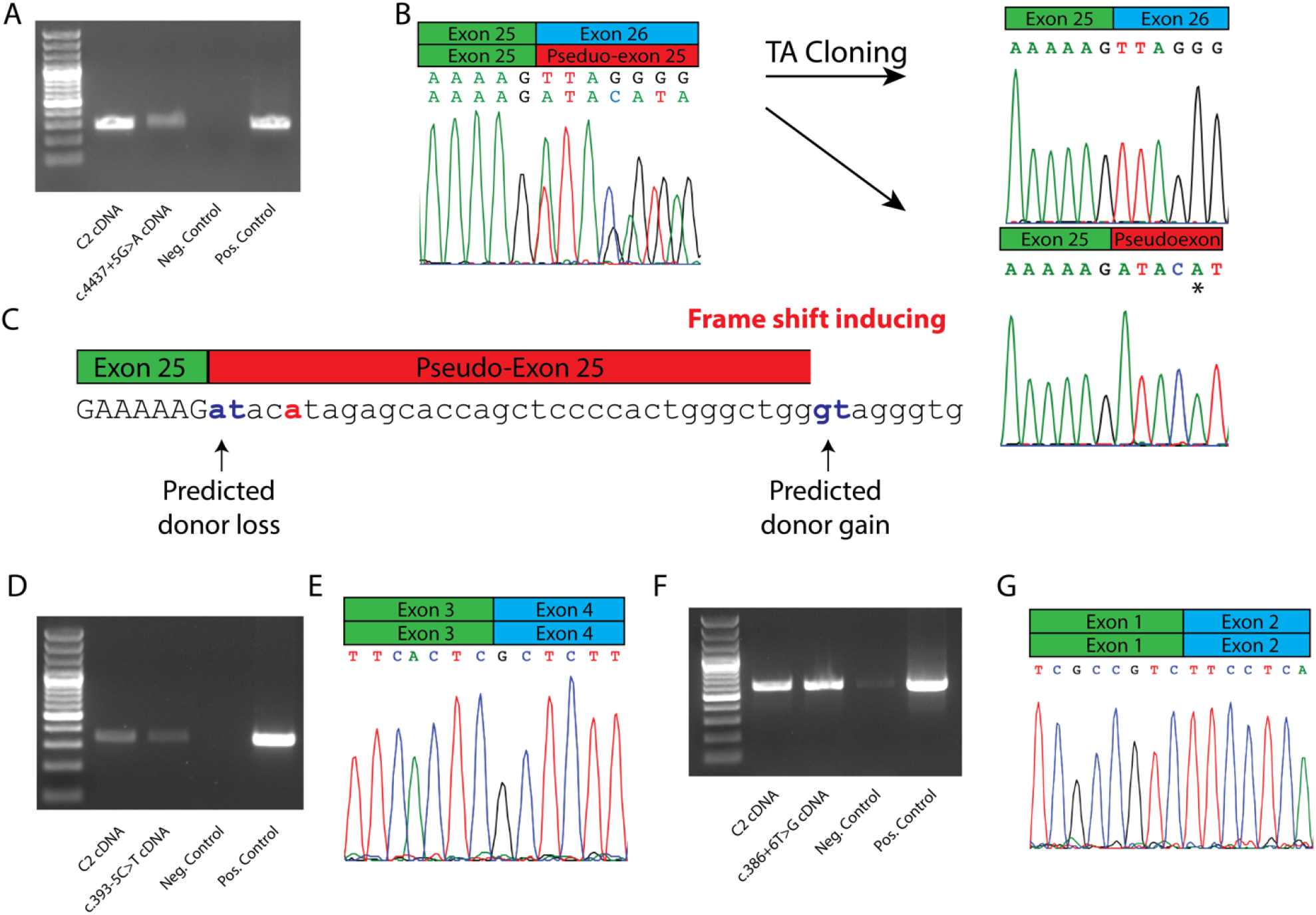
Variant functional analysis through CRISPR and iPSC-CMs. A) RT-PCR analysis of population control iPSC-CM (designated C2) vs *SCN5A* c.4437+5G>A iPSC-CM. Negative control indicates no PCR input, and positive control indicates *SCN5A* cDNA construct. B) Sanger trace of exon 25 junction shows normal splicing of WT allele and intron retention in the CRISPR-edited allele. * indicates variant DNA base from CRISPR edit. C) Schematic depiction of predicted splicing outcome of c.4437+5G>A, consistent with experimental findings. D) RT-PCR analysis of population control iPSC-CM (designated C2) vs *SCN5A* c.393-5C>T iPSC-CM. E) Sanger trace of exon 3/exon 4 junction shows WT splicing of both *SCN5A* alleles c.393-5C and c.393-5T. F) RT-PCR analysis of population control iPSC-CM (designated C2) vs *KCNQ1* c.386+6T>G iPSC-CM. G) Sanger trace of exon 1/exon 2 junction shows canonical splicing of both *KCNQ1* alleles c.386+6T>G.

### SpliceAI Scores of Affected Variants

Aggregate SpliceAI scores are shown in Table 1 (individual scores are provided in Supplementary Table 2). SpliceAI provides probabilities of losing or gaining a splice acceptor or donor site due to *cis*-genetic variation. Most variants were predicted to introduce and/or ablate a donor site as their major consequence, consistent with their position at the 3’ end of each exon (12/13). SpliceAI results were generally concordant with our experimental findings. For example, the *SCN5A* c.393-5C>T VUS was not predicted to alter splicing, matching the experimental findings (Figure 4C). 7/11 variants that disrupted splicing *in vitro* had SpliceAI scores >0.5. One exception is *KCNH2* c.2398+5G>T, which was not predicted to disrupt a splice site by SpliceAI, but was observed in 2 LQTS patients^13^ and caused exon skipping *in vitro*. SpliceAI also successfully identified the exonic cryptic donor splice site introduced by the LP variant *SCN5A* c.4719C>T (0.96 aggregate score).

### ACMG Reclassification of Putative Splice-altering Variants

Following functional assays, we applied the PS3 criteria to 9 variants and the BS3 to 1 variant. Computational predictions from SpliceAI were also included: 2 variants meeting BP4 and 3 meeting PP3 criteria. After integrating these functional and computational findings with the previous ACMG criteria calculated by Walsh et al.^13^ (Supplementary Table 3), we reclassified 9/12 VUS to LP, 1 VUS to LB, and 1 LP variant to P (Table 1).

## Discussion

### Splicing as Disease Mechanism

It is estimated that up to 10% of pathogenic variants may arise from aberrant splicing^8^. In addition to the widespread annotation of variants disrupting the canonical 2-bp splice sites as meeting the PVS1 criterion, the importance of splice-altering variants outside the canonical 2-bp splice sites has also been shown for Mendelian diseases^31,32^. While not all variation close to the exon/intron junction should be assumed to affect splicing, functional assays provide expedient evidence for prospective variant reclassification when observed. Although we did not use SpliceAI to help decide which variants to investigate, SpliceAI scores correlated well with functional assay results. This recently developed convolutional deep neural network appears to predict aberrant splicing with much higher accuracy than previous computational predictors^8^. Future *in vitro* splicing studies might therefore prioritize variants that are predicted to be splice-altering by this algorithm. Notably, the LQTS/BrS consortium dataset did not include any examples of deep intronic variation, commonly defined as >100 bp away from the exon/intron junction^33^.

### Functional Assays Reveal Aberrant Splicing

We applied functional studies to reclassify a set of VUS that have been observed in clinical cohorts of BrS and LQTS patients. The minigene remains a standard assay to address the fidelity of splicing and has been deployed in many applications^34,35^. Although a few reports of high-throughput adaptations of the minigene system have been published, these often remain restricted to a small genomic area^36^ or are limited by in-frame exon triplet cassettes^37^. A complementary tool is RT-PCR of primary patient tissue or a secondary tissue developed from patient-derived iPSCs when primary tissue is unavailable. Studying candidate arrhythmia variants in a cardiomyocyte context naturally provides more information than the *in vitro* minigene assays alone. For example, investigators found many more splicing aberrations in an iPSC-CM model than in minigene assays alone for the LQTS-associated splice variant c.1032G>A^28^. We observed aberrant splicing in 1/3 of our CRISPR edited iPSC-CM assays. We did not detect aberrant splicing for the *SCN5A* c.393-5C>T variant in iPSC-CMs, which was discordant with a previous study that showed exon skipping in a minigene assay^30^. As mentioned above, however, the use of non-canonical splice sites in minigene assays may have caused the previously observed exon skipping events. Further supporting our results is the high frequency of the related c.393-5C>A variant: an allele count of 45 in gnomAD, ∼8x higher than the 2.5e-5 cutoff that we and others have used^14^.

### Aberrant Splicing-associated Cardiac Morbidity

Aberrant splicing is increasingly recognized as a driver of cardiovascular disease. Most of such variation has been implicated in cardiomyopathy genes^31,32,38-40^. A recent functional genomics variant reclassification effort examined aberrant splicing from a set of 56 hypertrophic cardiomyopathy (HCM) probands with *MYBPC3* variants. The investigators performed RNA analysis of 9 such variants outside the canonical splice sites from venous blood or myocardial tissue, and were able to reclassify 6 variants (4 VUS -> LP, and 2 LP -> P)^40^. In a follow up study, the investigators used patient-derived iPSC-CMs followed by RNA-seq to identify 2 known and 1 novel deep intronic splice variants in HCM probands^38^. The investigators also found that rationally designed antisense oligonucleotides (ASOs) could correct the aberrant splicing of one of the three variants. In a similar study, investigators combined SpliceAI with cohort WGS to identify known splice-altering variants and a novel splice-altering variant in *MYBPC3* from studies of peripheral blood and RT-PCR^*39*^. A complementary study jointly investigated splice-altering variants in *LMNA*, a dilated cardiomyopathy (DCM) associated gene, and *MYBPC3* using computational tools to prioritize variants, and a high-throughput sequencing-based minigene platform^32^. The authors also extended this system to study non-canonical splice-altering variants in *TTN*^*31*^. Aberrant splicing in the cardiac arrhythmias is comparatively less studied. A seminal study identified a branch point-altering variant in *KCNH2* that elicited the LQTS phenotype in a multigenerational family using patient tissue and minigene assays^41^. More recently, a deep intronic splice-altering variant in *KCNH2* was identified in a genotype-negative LQTS family and functionally characterized using a similar CRISPR iPSC-CM framework as used in this study^42^.

### In silico Prediction Concordance

We found that SpliceAI agreed with our functional assays in most cases (Table 1). For example, our RT-PCR studies of the *SCN5A* c.393-5C>T variant agreed with the benign predictions of SpliceAI to have little effect. The *SCN5A* c.4719C>T variant was also strongly predicted by SpliceAI to introduce a new cryptic exonic donor site, consistent with the truncation event observed in our study. In contrast, we observed divergent outcomes with the *KCNQ1* c.386+6T>G variant, where we observed no splicing abnormality despite high predictions of disruption (0.96 aggregate score). SpliceAI also was highly predictive of disrupted splicing at the +5 position (4/5 variant studied had >0.5 aggregate SpliceAI score). The +5 position has frequently been invoked in disease and is particularly intolerant to variation in evolutionary studies^43^. Experimentally, a recent high-throughput study of non-canonical splice-altering *TTN* variants also found a large enrichment of splice-altering variants at this position^31^.

### Therapeutic Opportunities

Splice-altering variants comprise an eminently targetable class of genetic variation, with therapeutic modalities spanning ASOs, traditional small molecules, and emerging gene editing platforms. ASO technologies that leverage coding variation benefit by retaining pharmacokinetic and pharmacodynamic properties over a broad range of possible sequence targets^44^. Indeed, ‘N of 1’ drug discovery has already been implemented clinically^45^, and has even precipitated new regulatory processes in anticipation of increased translational opportunities^46^. Complementary mechanism-agnostic gene therapy approaches such as the ‘suppression-replacement’ strategy may be capable of treating a variety of variant classes in a specified gene, including those that alter splicing^47^. Altogether, the characterization of splice-altering variants may prioritize therapeutic opportunities in the genetic arrhythmias.

### Future Directions

Future studies can apply advances multiplexed assays to enable rapid high-throughput prospective evaluation of splice-perturbing variants. Further characterizing the fraction of splice-altering variants responsible for the Mendelian channelopathies may inspire future investigations in this area.

### Limitations

HEK293 cells do not capture the full environment of a native cardiomyocyte. Trans-acting proteins may also affect splicing outcomes in native cardiomyocytes. Although minigene assays offer clear functional readouts, the exact consequences of the reading frame in a human gene may remain cryptic due to the limited intronic window of the assay. Predicted termination codons beyond the 100 bp of cloned intronic DNA are therefore necessarily inferred from the observed reading frames. The variants in this study represent only a small fraction of the total quantity of possible splice site-adjacent variants in these three arrythmia-associated genes.

## Conclusions

We deployed functional assays to reclassify 11 variants near canonical splice sites observed in BrS and LQTS probands.

## Description of Supplementary Data

Supplementary data includes Supplementary Figures 1-3, and Supplementary Tables 1-3.

## Data Availability

All data used in this study is available upon reasonable request from the authors.

## Acknowledgements

Flow Cytometry experiments were performed in the VMC Flow Cytometry Shared Resource. The VMC Flow Cytometry Shared Resource is supported by the Vanderbilt Ingram Cancer Center (P30 CA68485) and the Vanderbilt Digestive Disease Research Center (DK058404). This research was funded by NIH grants T32GM007347 and AHA 907581 (MJO), the Heart Rhythm Society Clinical Research Award in Honor of Mark Josephson and Hein Wellens (Y.W.), AHA 830951 (Y.W.), K99 HG010904 (AMG), and R01 HL149826 (DMR).

## Author Contributions

Conceptualization: M.J.O., A.M.G., D.M.R.; Data curation: M.J.O., Y.W., L.D.H., D.W.M., A.M.G., D.M.R.; Formal Analysis: M.J.O., A.M.G., D.M.R.; Supervision: A.M.G., D.M.R.; Visualization: M.J.O., A.M.G., D.M.R.; Writing-original draft: M.J.O., A.M.G., D.M.R.; Writing-review & editing: M.J.O., Y.W., L.D.H., D.W.M., A.M.G., D.M.R.

## Ethics Declaration

All data generated from previous studies used in this analysis received IRB/REC approval.

## Conflicts of Interest

The authors have no disclosures.

## Notes

### Competing Interest Statement

The authors have declared no competing interest.

## References

1 Adler, A. et al. An International, Multicentered, Evidence-Based Reappraisal of Genes Reported to Cause Congenital Long QT Syndrome. Circulation 141, 418–428, doi:10.1161/circulationaha.119.043132 (2020).

2 Hosseini, S. M. et al. Reappraisal of Reported Genes for Sudden Arrhythmic Death: Evidence-Based Evaluation of Gene Validity for Brugada Syndrome. Circulation 138, 1195–1205, doi:10.1161/circulationaha.118.035070 (2018).

3 Schwartz, P. J. et al. Inherited cardiac arrhythmias. Nat Rev Dis Primers 6, 58, doi:10.1038/s41572-020-0188-7 (2020).

4 Glazer, A. M. et al. Arrhythmia variant associations and reclassifications in the eMERGE-III sequencing study. medRxiv, 2021.2003.2030.21254549, doi:10.1101/2021.03.30.21254549 (2021).

5 Parikh, V. N. Promise and Peril of Population Genomics for the Development of Genome-First Approaches in Mendelian Cardiovascular Disease. Circ Genom Precis Med 14, e002964, doi:10.1161/circgen.120.002964 (2021).

6 Richards, S. et al. Standards and guidelines for the interpretation of sequence variants: a joint consensus recommendation of the American College of Medical Genetics and Genomics and the Association for Molecular Pathology. Genet Med 17, 405–424, doi:10.1038/gim.2015.30 (2015).

7 Karczewski, K. J. et al. The mutational constraint spectrum quantified from variation in 141,456 humans. Nature 581, 434–443, doi:10.1038/s41586-020-2308-7 (2020).

8 Jaganathan, K. et al. Predicting Splicing from Primary Sequence with Deep Learning. Cell 176, 535–548.e524, doi:10.1016/j.cell.2018.12.015 (2019).

9 Sibley, C. R., Blazquez, L. & Ule, J. Lessons from non-canonical splicing. Nat Rev Genet 17, 407–421, doi:10.1038/nrg.2016.46 (2016).

10 Pertea, M., Lin, X. & Salzberg, S. L. GeneSplicer: a new computational method for splice site prediction. Nucleic Acids Res 29, 1185–1190, doi:10.1093/nar/29.5.1185 (2001).

11 Yeo, G. & Burge, C. B. Maximum entropy modeling of short sequence motifs with applications to RNA splicing signals. J Comput Biol 11, 377–394, doi:10.1089/1066527041410418 (2004).

12 Hanses, U. et al. Intronic CRISPR Repair in a Preclinical Model of Noonan Syndrome-Associated Cardiomyopathy. Circulation 142, 1059–1076, doi:10.1161/circulationaha.119.044794 (2020).

13 Walsh, R. et al. Enhancing rare variant interpretation in inherited arrhythmias through quantitative analysis of consortium disease cohorts and population controls. Genet Med 23, 47–58, doi:10.1038/s41436-020-00946-5 (2021).

14 Whiffin, N. et al. Using high-resolution variant frequencies to empower clinical genome interpretation. Genet Med 19, 1151–1158, doi:10.1038/gim.2017.26 (2017).

15 Schneider, C. A., Rasband, W. S. & Eliceiri, K. W. NIH Image to ImageJ: 25 years of image analysis. Nat Methods 9, 671–675, doi:10.1038/nmeth.2089 (2012).

16 Concordet, J. P. & Haeussler, M. CRISPOR: intuitive guide selection for CRISPR/Cas9 genome editing experiments and screens. Nucleic Acids Res 46, W242–w245, doi:10.1093/nar/gky354 (2018).

17 Ran, F. A. et al. Genome engineering using the CRISPR-Cas9 system. Nat Protoc 8, 2281–2308, doi:10.1038/nprot.2013.143 (2013).

18 Wada, Y. et al. Common Ancestry-specific Ion Channel Variants Predispose to Drug-induced Arrhythmias. Circulation, doi:10.1161/circulationaha.121.054883 (2022).

19 Burridge, P. W. et al. Chemically defined generation of human cardiomyocytes. Nat Methods 11, 855–860, doi:10.1038/nmeth.2999 (2014).

20 Blakes, A. J. M. et al. A systematic analysis of splicing variants identifies new diagnoses in the 100,000 Genomes Project. medRxiv, 2022.2001.2028.22270002, doi:10.1101/2022.01.28.22270002 (2022).

21 Bastarache, L. et al. Phenotype risk scores identify patients with unrecognized Mendelian disease patterns. Science 359, 1233–1239, doi:10.1126/science.aal4043 (2018).

22 Valenzuela-Palomo, A. et al. Splicing predictions, minigene analyses, and ACMG-AMP clinical classification of 42 germline PALB2 splice-site variants. J Pathol, doi:10.1002/path.5839 (2021).

23 Fraile-Bethencourt, E. et al. Functional classification of DNA variants by hybrid minigenes: Identification of 30 spliceogenic variants of BRCA2 exons 17 and 18. PLoS Genet 13, e1006691, doi:10.1371/journal.pgen.1006691 (2017).

24 Kleinberger, J., Maloney, K. A., Pollin, T. I. & Jeng, L. J. in Genet Med Vol. 18 1165 (2016).

25 Laššuthová, P. et al. Clinical, in silico, and experimental evidence for pathogenicity of two novel splice site mutations in the SH3TC2 gene. J Neurogenet 26, 413–420, doi:10.3109/01677063.2012.711398 (2012).

26 Cho, S. Y. et al. Novel PPOX exonic mutation inducing aberrant splicing in a patient with homozygous variegate porphyria. Clin Chim Acta 512, 117–120, doi:10.1016/j.cca.2020.10.033 (2021).

27 Bardai, A. et al. Sudden cardiac arrest associated with use of a non-cardiac drug that reduces cardiac excitability: evidence from bench, bedside, and community. Eur Heart J 34, 1506–1516, doi:10.1093/eurheartj/eht054 (2013).

28 Wuriyanghai, Y. et al. Complex aberrant splicing in the induced pluripotent stem cell-derived cardiomyocytes from a patient with long QT syndrome carrying KCNQ1-A344Aspl mutation. Heart Rhythm 15, 1566–1574, doi:10.1016/j.hrthm.2018.05.028 (2018).

29 Burset, M., Seledtsov, I. A. & Solovyev, V. V. Analysis of canonical and non-canonical splice sites in mammalian genomes. Nucleic Acids Res 28, 4364–4375, doi:10.1093/nar/28.21.4364 (2000).

30 Frisso, G. et al. Functional Studies and In Silico Analyses to Evaluate Non-Coding Variants in Inherited Cardiomyopathies. Int J Mol Sci 17, doi:10.3390/ijms17111883 (2016).

31 Patel, P. N. et al. Contribution of Noncanonical Splice Variants to TTN Truncating Variant Cardiomyopathy. Circ Genom Precis Med 14, e003389, doi:10.1161/circgen.121.003389 (2021).

32 Ito, K. et al. Identification of pathogenic gene mutations in LMNA and MYBPC3 that alter RNA splicing. Proc Natl Acad Sci U S A 114, 7689–7694, doi:10.1073/pnas.1707741114 (2017).

33 Vaz-Drago, R., Custódio, N. & Carmo-Fonseca, M. Deep intronic mutations and human disease. Hum Genet 136, 1093–1111, doi:10.1007/s00439-017-1809-4 (2017).

34 Gaildrat, P. et al. Use of splicing reporter minigene assay to evaluate the effect on splicing of unclassified genetic variants. Methods Mol Biol 653, 249–257, doi:10.1007/978-1-60761-759-4_15 (2010).

35 Gao, D. et al. A deep learning approach to identify gene targets of a therapeutic for human splicing disorders. Nat Commun 12, 3332, doi:10.1038/s41467-021-23663-2 (2021).

36 Gergics, P. et al. High-throughput splicing assays identify missense and silent splice-disruptive POU1F1 variants underlying pituitary hormone deficiency. Am J Hum Genet, doi:10.1016/j.ajhg.2021.06.013 (2021).

37 Cheung, R. et al. A Multiplexed Assay for Exon Recognition Reveals that an Unappreciated Fraction of Rare Genetic Variants Cause Large-Effect Splicing Disruptions. Mol Cell 73, 183–194.e188, doi:10.1016/j.molcel.2018.10.037 (2019).

38 Holliday, M. et al. Transcriptome Sequencing of Patients With Hypertrophic Cardiomyopathy Reveals Novel Splice-Altering Variants in MYBPC3. Circ Genom Precis Med 14, e003202, doi:10.1161/circgen.120.003202 (2021).

39 Lopes, L. R. et al. Cryptic Splice-Altering Variants in MYBPC3 Are a Prevalent Cause of Hypertrophic Cardiomyopathy. Circ Genom Precis Med 13, e002905, doi:10.1161/circgen.120.002905 (2020).

40 Singer, E. S., Ingles, J., Semsarian, C. & Bagnall, R. D. Key Value of RNA Analysis of MYBPC3 Splice-Site Variants in Hypertrophic Cardiomyopathy. Circ Genom Precis Med 12, e002368, doi:10.1161/circgen.118.002368 (2019).

41 Crotti, L. et al. A KCNH2 branch point mutation causing aberrant splicing contributes to an explanation of genotype-negative long QT syndrome. Heart Rhythm 6, 212–218, doi:10.1016/j.hrthm.2008.10.044 (2009).

42 Tobert, K. E. et al. Genome Sequencing in a Genetically Elusive Multi-Generational Long QT Syndrome Pedigree Identifies a Novel LQT2-Causative Deeply Intronic KCNH2 Variant. Heart Rhythm, doi:10.1016/j.hrthm.2022.02.004 (2022).

43 Lord, J. et al. Pathogenicity and selective constraint on variation near splice sites. Genome Res 29, 159–170, doi:10.1101/gr.238444.118 (2019).

44 Crooke, S. T., Baker, B. F., Crooke, R. M. & Liang, X. H. Antisense technology: an overview and prospectus. Nat Rev Drug Discov 20, 427–453, doi:10.1038/s41573-021-00162-z (2021).

45 Kim, J. et al. Patient-Customized Oligonucleotide Therapy for a Rare Genetic Disease. N Engl J Med 381, 1644–1652, doi:10.1056/NEJMoa1813279 (2019).

46 Brierley, J., Aylett, S. & Archard, D. Framework for “N-of-1” Experimental Therapies. N Engl J Med 382, e7, doi:10.1056/NEJMc1915778 (2020).

47 Dotzler, S. M. et al. Suppression-Replacement KCNQ1 Gene Therapy for Type 1 Long QT Syndrome. Circulation, doi:10.1161/circulationaha.120.051836 (2021).

