## Supplementary Data for "Functional Assays Reclassify Suspected Splice-Altering Variants of Uncertain Significance in Mendelian Channelopathies"

### Supplement

Table of Contents

**Table S1:** Primers used in this study.
**Table S2:** SpliceAI Predictions by Variant.
**Table S3:** Previous ACMG Reclassification with Population Data.

**Figure S1:** Inconclusive minigene results for 2 *SCN5A* VUS.
**Figure S2**: *KCNH2* Minigene Assay Gels and Sanger Traces.
**Figure S3:** *KCNQ1* Minigene Assay Gels and Sanger Traces.

**Table S1. Primers used in this study.**

| Primer | Name | Sequence |
|---|---|---|
| *SCN5A* c.1890G>A (for) | mo51 | GCCGGTCGACTGCTCTGAGAAGTTTAGCTGAGG |
| *SCN5A* c.1890G>A (rev) | mo52 | GCCGGCGGCCGCCACACAGTAGGTGCTCAACAA |
| *SCN5A* c.4299+6T>C (for) | mo8 | GCCGGTCGACCCCTCTTTCCCACAGAATGG |
| *SCN5A* c.4299+6T>C (rev) | mo9 | GCCGGCGGCCGCGAAGAGGACCATCCCCAACA |
| *SCN5A* c.4299+G>C (for) | mo8 | GCCGGTCGACCCCTCTTTCCCACAGAATGG |
| *SCN5A* c.4299+G>C (rev) | mo9 | GCCGGCGGCCGCGAAGAGGACCATCCCCAACA |
| *SCN5A* c.4719C>T (for) | mo11 | GCCGGTCGACGAGTGGGAGGGTGGGTGG |
| *SCN5A* c.4719C>T (rev) | mo12 | GCCGGCGGCCGCAAGAAGCTAGGGTTGTACATGGC |
| *KCNH2* c.1128G>A (for) | mo15 | GCCGGTCGACCCCATGGACTCCTGACTGTG |
| *KCNH2* c.1128G>A (rev) | mo16 | GCCGGCGGCCGCTCACTGGCTAGCCCCCTC |
| *KCNH2* c.2145G>A (for) | mo18 | GCCGGTCGACGGCATGTCGACGCTGAGAC |
| *KCNH2* c.2145G>A (rev) | mo19 | GCCGGCGGCCGCCGTGCCTCCAGGGTCCTTAC |
| *KCNH2* c.2398+5G>T (for) | mo21 | GCCGGTCGACTCTCCCAGGCCTGGAGGT |
| *KCNH2* c.2398+5G>T (rev) | mo22 | GCCGGCGGCCGCGAGCCCAGTGACCCTGCA |
| *KCNQ1* c.477+5G>A (for) | mo24 | GCCGGTCGACCACTGTCTGTTCCTACCTGGG |
| *KCNQ1* c.477+5G>A (rev) | mo25 | GCCGGCGGCCGCGCAAGGGGCTGGGAGTG |
| *KCNQ1* c.683+5G>A (for) | mo27 | GCCGGTCGACGACCCGTCTGACCAGCAAG |
| *KCNQ1* c.683+5G>A (rev) | mo28 | GCCGGCGGCCGCGGCAGTGGGCACCAAGG |
| *KCNQ1* c.1032G>A (for) | mo31 | GCCGGTCGACGTCTCTTGCCGGCCTCTC |
| *KCNQ1* c.1032G>A (rev) | mo32 | GCCGGCGGCCGCGAACGTAAGTGGGTCTGCTCA |
| *SCN5A* c.1890G>A (for) | mo51 | GCCGGTCGACTGCTCTGAGAAGTTTAGCTGAGG |
| *SCN5A* c.1890G>A (rev) | mo52 | GCCGGCGGCCGCCACACAGTAGGTGCTCAACAA |
| *SCN5A* c.4299+6T>C (for) | mo8 | GCCGGTCGACCCCTCTTTCCCACAGAATGG |
| *SCN5A* c.4299+6T>C (rev) | mo9 | GCCGGCGGCCGCGAAGAGGACCATCCCCAACA |
| *SCN5A* c.4299+G>C (for) | mo8 | GCCGGTCGACCCCTCTTTCCCACAGAATGG |
| *SCN5A* c.4299+G>C (rev) | mo9 | GCCGGCGGCCGCGAAGAGGACCATCCCCAACA |
| *SCN5A* c.4719C>T (for) | mo11 | GCCGGTCGACGAGTGGGAGGGTGGGTGG |
| *SCN5A* c.4719C>T (rev) | mo12 | GCCGGCGGCCGCAAGAAGCTAGGGTTGTACATGGC |
| *SCN5A* c.1890G>A | mo7 | CCCGCCAGACACAGTGAGCCAGCCC |
| *SCN5A* c.4299+6T>C | mo14 | CAGGGGGGTAGGCTGCCACAGTGGC |
| *SCN5A* c.4299+G>C | mo2 | GGACTCCAGGGGCGTAGGTTGCCAC |
| *SCN5A* c.4719C>T | mo13 | TGGCCATCTTCACAGGTGAGTGTATTGTCAAGCT |
| *KCNH2* c.1128G>A | mo17 | AGAAGGTCACCCAAGTAGGCGCCCAGC |
| *KCNH2* c.2145G>A | mo20 | CATCGACATGAACGCAGTGAGGCCACCAGAG |
| *KCNH2* c.2398+5G>T | mo23 | GTGGCCATCCTGGGTATTGGGTGGGGG |
| *KCNQ1* c.477+5G>A | mo26 | TCTCTTCTGGATGGTACATAGCATCTGAGGGCATG |
| *KCNQ1* c.683+5G>A | mo29 | CGGCCATCAGGTGCATCTGTGCCACAAGC |
| *KCNQ1* c.1032G>A | mo36 | GCGCTCCCAGCAGTAGGTGCCCC |
| *KCNQ1* c.1032G>T | mo35 | GCGCTCCCAGCTGTAGGTGCCCC |
| *KCNQ1* c.1032+5G>A | mo32 | CCCAGCGGTAGATGCCCCGTGGG |
| *SCN5A* c.1890G>A | mo7 | CCCGCCAGACACAGTGAGCCAGCCC |
| *SCN5A* c.4299+6T>C | mo14 | CAGGGGGGTAGGCTGCCACAGTGGC |
| *SCN5A* c.4299+G>C | mo2 | GGACTCCAGGGGCGTAGGTTGCCAC |
| *SCN5A* c.393-5C>T (for) | mo42 | CACCGAGCATGTTGAAGAGCGTGCG |
| *SCN5A* c.393-5C>T (rev) | mo43 | AAACCGCACGCTCTTCAACATGCTC |
| *SCN5A* c.393-5C>T (repair template) | mo61 | ATGACCCCCAGGGGCAGGACCCCCCC<br>AGCGGTGGTGGCGTGGCCGGCCCATGCTGCT<br>CAGCTTTCCTTGACCACATGCACGCTCTTCAAC<br>ATGCTCATCATGTGCACCATCCTCA<br>CCAACTGCGTGTTCATGGCCCA<br>GCACGACCCTCCA |
| *SCN5A* c.393-5C>T sequencing (for) | mo5 | GCCGGTCGACTTGTCTGGTAGCACTGGC |

| | | |
|---|---|---|
| *SCN5A* c.393-5C>T sequencing (rev) | mo6 | GCCGGCGGCCGCCCTGGGCCTGGACACAAG |
| *SCN5A* c.4437+5G>A (for) | mo62 | CACC*G*AGCTCCCCACTGGGCTGGGT |
| *SCN5A* c.4437+5G>A (rev) | mo63 | AAACACCCAGCCCAGTGGGGAGCT*C* |
| *SCN5A* c.4437+5G>A (repair template) | mo64 | TGGAGCCTGAGTGGCCCCTCAA<br>TCCCCTGGCACCCGGCCCCACCGTACCCAGC<br>CCAGTGGGGAGCTGGTGCTCTATGTATCTTTTT<br>CTTCTGTTGGTTGAAGTTGTCAATG<br>ATGACACCAATAAAGAGGTT<br>CAGGGTGAAGAAAGACCCA |
| *SCN5A* c.4437+5G>A sequencing (for) | mo8 | GCCGGTCGACCCCTCTTTCCCACAGAATGG |
| *SCN5A* c.4437+5G>A sequencing (rev) | mo9 | GCCGGCGGCCGCGAAGAGGACCATCCCCAACA |
| *KCNQ1* c.386+6T>G (for) | mo55 | CACCGTGAGTATCGCCACCGGCGA |
| *KCNQ1* c.386+6T>G (rev) | mo56 | AAACTCGCCGGTGGCGATACTCAC |
| *KCNQ1* c.386+6T>G (repair template) | mo57 | GCGCTGGGACAGAGCTCCCCCAC<br>ACCAGCTCTCAGGAAGCACCTTCG<br>TGCCGGCGGTCGCCGGTGGCGATC<br>CTCACACGGCGAAGTGGTAAACGAAGC<br>ATTTCCAGCCGGTGGGACGCTCGAG<br>GAAGTTGTAGACGCGGCCCTGGACGTG |
| *KCNQ1* c.386+6T>G sequencing (for) | mo58 | ATCTACAGCACGCGCCG |
| *KCNQ1* c.386+6T>G sequencing (rev) | mo59 | CTCCTCTGCTCCGGGGT |
| *SCN5A* c.393-5C>T (for) | mo42 | CACCGAGCATGTTGAAGAGCGTGCG |
| *SCN5A* c.393-5C>T (rev) | mo43 | AAACCGCACGCTCTTCAACATGCTC |
| *SCN5A* c.393-5C>T (repair template) | mo61 | ATGACCCCCAGGGGCAGGACCCCCCC<br>AGCGGTGGTGGCGTGGCCGGCCCATGCTGCT<br>CAGCTTTCCTTGACCACATGCACGCTCTTCAAC<br>ATGCTCATCATGTGCACCATCCTCA<br>CCAACTGCGTGTTCATGGCCCA<br>GCACGACCCTCCA |
| *SCN5A* c.393-5C>T sequencing (for) | mo5 | GCCGGTCGACTTGTCTGGTAGCACTGGC |
| *SCN5A* c.393-5C>T sequencing (rev) | mo6 | GCCGGCGGCCGCCCTGGGCCTGGACACAAG |
| *SCN5A* c.4437+5G>A (for) | mo62 | CACC*G*AGCTCCCCACTGGGCTGGGT |
| *SCN5A* c.4437+5G>A (rev) | mo63 | AAACACCCAGCCCAGTGGGGAGCT*C* |
| *SCN5A* c.4437+5G>A (repair template) | mo64 | TGGAGCCTGAGTGGCCCCTCAA<br>TCCCCTGGCACCCGGCCCCACCGTACCCAGC<br>CCAGTGGGGAGCTGGTGCTCTATGTATCTTTTT<br>CTTCTGTTGGTTGAAGTTGTCAATG<br>ATGACACCAATAAAGAGGTT<br>CAGGGTGAAGAAAGACCCA |
| *SCN5A* c.4437+5G>A sequencing (for) | mo8 | GCCGGTCGACCCCTCTTTCCCACAGAATGG |
| *SCN5A* c.4437+5G>A sequencing (rev) | mo9 | GCCGGCGGCCGCGAAGAGGACCATCCCCAACA |
| *KCNQ1* c.386+6T>G (for) | mo55 | CACCGTGAGTATCGCCACCGGCGA |
| pET01 Sequencing (for) | mo37 | GGATTCTTCTACACACCC |
| pET01 Sequencing (alt for) | mo102 | CCTTTGTGGTTCTCACTTGGT |
| pET01 Sequencing (rev) | ag489 | GGGCCACCTCCAGTGC |
| pET01 RT | mo38 | GATCCACGATGC |

**Table S2. Previous ACMG Reclassification with Population Data[13].**

| Gene | CDS | Cases-Europe | Cases-Japan | ACMG Pre | ACMG Post | Reclassification |
|---|---|---|---|---|---|---|
| *SCN5A* | c.393-5C>T | 1 | 0 | PM2 | PM2 | VUS → VUS |
| *SCN5A* | c.1890G>A | 1 | 0 | PM2 | PM2 | VUS → VUS |
| *SCN5A* | c.4299+6T>C | 1 | 0 | PM2 | PM2 | VUS → VUS |
| *SCN5A* | c.4299G>C | 1 | 0 | PM2 | PM2 | VUS → VUS |
| *SCN5A* | c.4437+5G>A | 2 | 0 | PM2 | PM2 | VUS → VUS |
| *SCN5A* | c.4719C>T | 2 | 0 | PM2/PS3 | PM2, PS3, PS4_strong | LP → LP |
| *KCNH2* | c.1128G>A | 2 | 0 | PM2 | PM2 | VUS → VUS |
| *KCNH2* | c.2145G>A | 2 | 0 | PM2 | PM2 | VUS → VUS |
| *KCNH2* | c.2398+5G>T | 1 | 0 | PM2 | PM2 | VUS → VUS |
| *KCNQ1* | c.386+6T>G | 1 | 0 | PM2 | PM2 | VUS → VUS |
| *KCNQ1* | c.477+5G>A | 7 | 0 | PM2 | PM2, PS4_strong | VUS → LP |
| *KCNQ1* | c.683+5G>A | 3 | 0 | PM2 | PM2, PS4_moderate | VUS → VUS |
| *KCNQ1* | c.1032+5G>A | 2 | 0 | PM2 | PM2 | VUS → VUS |
| *KCNQ1* | c.1032G>T | 1 | 0 | PM2 | PM2 | VUS → VUS |
| *KCNQ1* | c.1032G>A | 9 | 20 | PM2 | PM2, PS4_strong | VUS → LP |

**Table S3. SpliceAI Predictions by Variant.**

| Gene | Variant | AG | AL | DG | DL | Aggregate |
|---|---|---|---|---|---|---|
| *SCN5A* | c.393-5C>T | 0 | 0.14 | 0 | 0.08 | 0.21 |
| *SCN5A* | c.1890G>A | 0 | 0 | 0.14 | 0.67 | 0.72 |
| *SCN5A* | c.4299+6T>C | 0 | 0.08 | 0.02 | 0.72 | 0.75 |
| *SCN5A* | c.4299G>C | 0 | 0.06 | 0.03 | 0.78 | 0.80 |
| *SCN5A* | c.4437+5G>A | 0.05 | 0.01 | 0.5 | 0.34 | 0.69 |
| *SCN5A* | c.4719C>T | 0 | 0.15 | 0.88 | 0.57 | 0.96 |
| *KCNH2* | c.1128G>A | 0 | 0.06 | 0.24 | 0.07 | 0.34 |
| *KCNH2* | c.2145G>A | 0.01 | 0.04 | 0.21 | 0 | 0.25 |
| *KCNH2* | c.2398+5G>T | 0 | 0.03 | 0.30 | 0.19 | 0.45 |
| *KCNQ1* | c.386+6T>G | 0.05 | 0 | 0.63 | 0.89 | 0.96 |
| *KCNQ1* | c.477+5G>A | 0 | 0.10 | 0.64 | 0.23 | 0.75 |
| *KCNQ1* | c.683+5G>A | 0 | 0.06 | 0.21 | 0.04 | 0.29 |
| *KCNQ1* | c.1032+5G>A | 0 | 0.01 | 0.81 | 0.75 | 0.95 |
| *KCNQ1* | c.1032G>T | 0 | 0.01 | 0.5 | 0.48 | 0.74 |
| *KCNQ1* | c.1032G>A | 0 | 0 | 0.49 | 0.38 | 0.68 |

Acceptor Gain (AG), Acceptor Loss (AL), Donor Gain (DG), Donor Loss (DL).

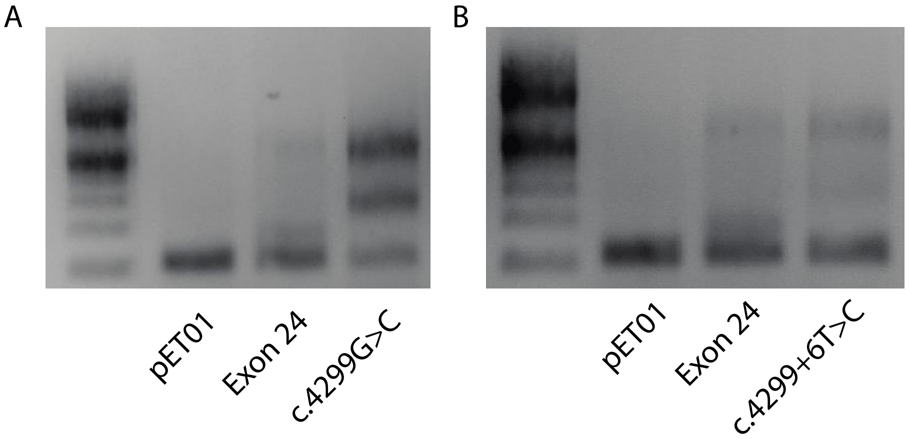


**Supplemental Figure 1. Inconclusive minigene results for 2 *SCN5A* VUS.**

A) Minigene results for VUS *SCN5A* c.4299G>C. WT exon 24 shows PSI <35%; however, 2 bands are present in *SCN5A* c.4299G>C lane corresponding to aberrant intron retention compared to reference sequence, as confirmed by Sanger sequencing.

B) Minigene results for VUS *SCN5A* c.4299+6T>C. Low WT PSI. Sequence of bands present in c.4299+6T>C was indeterminate.

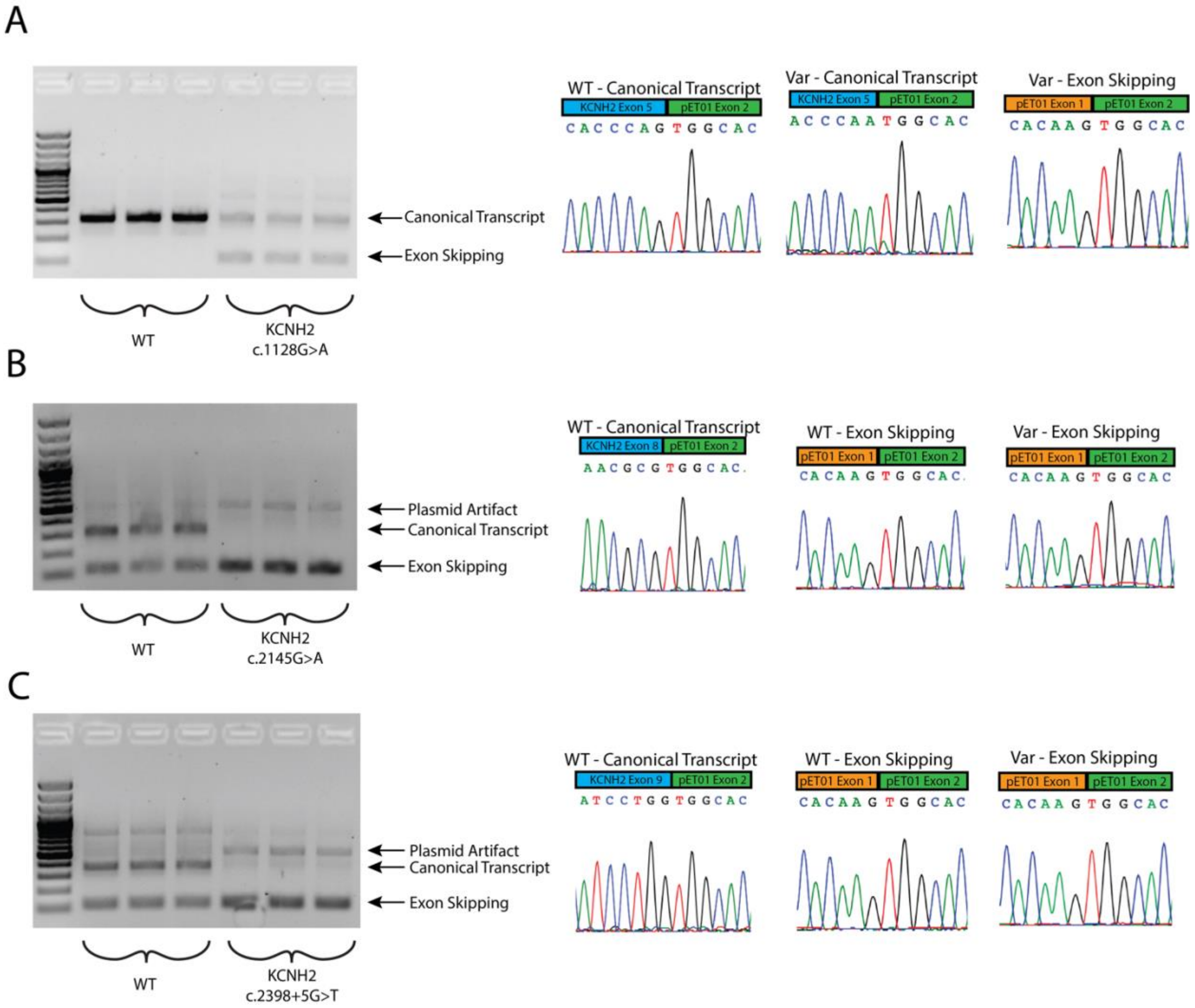


**Supplemental Figure S2. *KCNH2* Minigene Assay Gels and Sanger Traces.**

A) Gel analysis of RT-PCR products of *KCNH2* c.1128G>A. Two major products in the variant samples correspond to the WT transcript and the exon skipping product.

B) Gel analysis of RT-PCR products of *KCNH2* c.2145G>A. While the WT samples show exon skipping, the introduction of the variant leads to decreased WT PSI and an increase in exon skipping.

C) Gel analysis of RT-PCR products of *KCNH2* c.2398+5G>T. Introduction of variant leads to exon skipping.

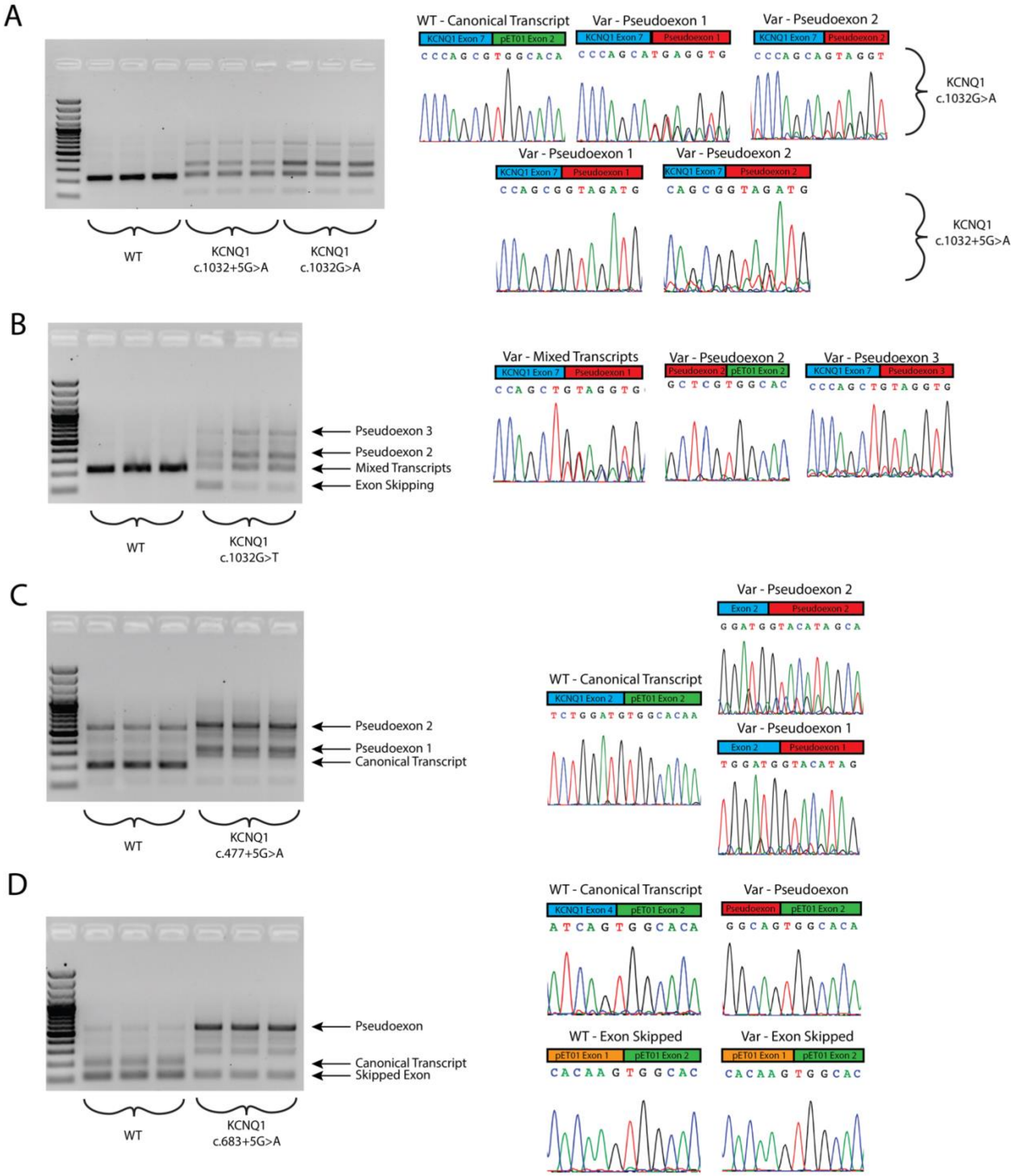


**Supplemental Figure S3.** ***KCNQ1*** **Minigene Assay Gels and Sanger Traces.**

A) Gel analysis of *KCNQ1* c.1032G>A and c.1032+5G>A variants. Each variant introduces multiple pseudoexons profiled by Sanger sequencing.

B) The c.1032G>T VUS introduces similar pseudoexons, while inducing a higher degree of exon skipping than that observed in c.1032G>A.

C) Gel electrophoresis of RT-PCR products show that the LP variant c.477+5G>A was observed to introduce at least 2 discrete pseudoexons and ablate WT splicing.

D) The VUS c.683+5G>A variant introduces a major pseudoexon while also introducing a degree of exon skipping.
